# Modeling healthy proteomic profiles for anomaly detection using subspace learning based one-class classification

**DOI:** 10.64898/2026.04.30.722013

**Authors:** Fahad Sohrab, Atul Kumar, Virpi Ahola, Andrew Magis, Ville Hautamäki, Merja Heinäniemi, Sui Huang

**Author notes:** Corresponding author(s). E-mail(s); **Correspondence and requests for materials** Correspondence and requests for materials should be addressed to F.S. S.H. ad M.H. contributed equally as co-senior authors. Contributing authors.

## Abstract

High-throughput plasma proteomics provides sensitive and scalable measurements of thousands of systemic protein profiles from minimally invasive blood samples, creating new opportunities for disease detection and population-scale health monitoring. However, robust statistical modeling remains challenging due to high dimensionality, limited availability and high diversity of diseased samples, resulting in class imbalance in clinical cohorts. Here, we present a subspace One-Class Classification (OCC) framework for proteomics-driven anomaly detection that models healthy proteomic profiles as a reference distribution. To address the limitations of conventional hyperparameter tuning in severely imbalanced data settings, we introduce a fully data-driven parameter estimation strategy that infers all model parameters directly from intrinsic properties of the healthy training data, without using any disease labels. Using plasma proteomics data generated with Olink, we evaluate a family of subspace and graph-embedded subspace extensions of Support Vector Data Description, in which all models operate on learned low-dimensional representations rather than the original feature space. Models are trained exclusively on a healthy reference cohort and evaluated on heterogeneous disease conditions, including multiple cancer types and an independent COVID-19 cohort, with all disease samples withheld from training to enable unbiased assessment of cross-disease generalization. Across disease contexts, the evaluated one-class models yield stable and balanced detection performance, demonstrating that learning structured low-dimensional representations of healthy proteomic variation captures intrinsic biological organization that generalizes across disease-specific perturbations. These results establish healthy-profile-based, subspace one-class learning as a robust and disease-agnostic framework for screening in high-dimensional plasma proteomics.

## 1 Introduction

Plasma proteomics, the measurement of thousands of circulating proteins including cytokines, hormones and intracellular leakage products, enables characterization of physiological and pathological states using minimally invasive blood samples [1–3]. Large-scale population studies have further demonstrated that plasma proteomic profiles can detect subclinical early stages or predict future risk of multiple diseases, highlighting the broad clinical potential of circulating proteome measurements [4, 5]. One leading technology for such high-dimensional profiling of protein abundances in plasma samples is Olink’s Proximity Extension Assay (PEA) that affords reproducible antibody-based quantification of hundreds to thousands of proteins in small sample volumes with high sensitivity and specificity [6–8]. These large-scale proteomic profiles capture coordinated biological responses across various domains, including immune, inflammatory and metabolic pathways, offering possibilities for disease biomarker discovery, risk stratification and population-scale screening [9, 10]. Despite these technological advances, extracting robust and generalizable disease-related signals from high-dimensional plasma proteomics data remains a major methodological challenge. A central difficulty in proteomics-based disease modeling lies in the statistical structure of clinical datasets [11]. As with all omics technologies, proteomic studies typically measure far more protein features than available samples, resulting in high dimensionality of features relative to sample number. This imbalance is further compounded by the limited availability, biological heterogeneity, and incomplete characterization of disease cohorts (samples with “ground truth” labels, such as otherwise verified diagnosis), whereas healthy reference samples are typically more abundant, stable, and better defined, although heterogeneity in this group can pose problems [12, 13]. Under these conditions, conventional binary or multiclass classification approaches are prone to overfitting and to unstable decision boundaries. Moreover, the dependence on balanced and representative data for training limits their utility for early-stage disease class discovery (as opposed to disease class detection), i.e. in the disease-agnostic evaluation of populations when disease-defining features are not known.

A key practical challenge for the clinical deployment of Olink and similar high-throughput proteomics technologies is that the disease relevance of most measured proteins remains largely unknown. Unlike established diagnostic laboratory variables, such as fasting serum glucose or cholesterol, for which well-defined physiological reference ranges exist, most proteins measured in large-scale plasma proteomic panels lack clearly defined normal abundance ranges. Consequently, it is often unclear what constitutes an “abnormal” value independent of the specific disease it might indicate. In clinical diagnostics, the interpretation of biomarkers typically begins with the definition of normal physiological variation, from which deviations can then be evaluated in the context of specific diseases. For large-scale proteomics, however, such reference boundaries are largely undefined. Addressing this gap requires first establishing the structure and limits of healthy proteomic variation. Defining this healthy proteomic state is therefore a prerequisite for identifying abnormal profiles of unknown characteristics in population-scale plasma proteomics and for enabling downstream disease-specific interpretation [14].

One-Class Classification (OCC) offers a formal path towards answering this pivotal clinical question by providing an alternative paradigm to traditional class discovery or class detection. Here, only a single well-characterized reference class is modeled, which most often represents the healthy state. This approach provides a principled way to define the boundaries of physiologically normal plasma proteomic profiles in a healthy cohort. Methods such as Support Vector Data Description (SVDD) learn a compact boundary around the normal samples and identify deviations as potential anomalies [15, 16]. This paradigm is particularly appealing for clinical proteomics, where diseased samples are often scarce, heterogeneous, or unavailable at training time.

Plasma proteomic profiles are inherently high-dimensional but exhibit substantial correlation structure, with many proteins showing coordinated variation due to shared biological pathways, leading to redundancy across features. As a result, feature-space distance and boundary-based descriptions degrade rapidly with increasing dimensionality, leading to unstable decision surfaces and reduced generalization [17]. These effects are particularly pronounced within the new systems biology and omics profiling setting where sample sizes are small relative to the number of features, which further amplify instability in distance-based learning [18].

Subspace-based OCC approaches address these limitations by coupling anomaly detection with representation learning. Instead of modeling samples in the original high-dimensional space, these methods learn a low-dimensional subspace in which variation becomes more compact and structured, making it well suited to capture coordinated patterns of protein expression that reflect underlying biological processes in the healthy cohort. Subspace Support Vector Data Description (SSVDD) and its extensions jointly optimize dimensionality reduction and data description, suppressing redundant variation while preserving coordinated protein responses that reflect underlying regulatory and signaling processes [19, 20]. This formulation is particularly well suited to plasma proteomics, where biological pathways give rise to low-dimensional manifolds embedded within high-dimensional measurement spaces. Similar low-dimensional manifold structures have been reported across diverse high-throughput biological modalities, including transcriptomics and proteomics, reflecting coordinated regulatory programs [21, 22].

In this study, we investigate healthy-profile-based one-class modeling as a disease-agnostic strategy for proteomics-driven disease detection, with subspace learning as an integral component of the modeling framework. Using plasma proteomics data generated on the Olink platform we systematically evaluate various subspace learning based OCC methods under both linear and non-linear formulations. All models operate exclusively in learned low-dimensional representations and are trained solely on a healthy reference cohort. Evaluation, the detection of “anomalies”, is then performed on heterogeneous disease conditions, including multiple cancer types and an independent COVID-19 cohort, with all disease samples withheld from training to enable unbiased assessment of cross-disease generalization.

The contributions of this work are threefold. First, we formulate the clinical problem of identifying abnormal plasma proteomic profiles as a strictly healthy-cohort-trained OCC problem, reflecting the fundamental idea of the existence of a physiologically normal space of plasma protein abundance configurations and the practical reality that a comprehensive and stable representation of disease spaces in high-dimensional proteomics is unlikely to ever be complete. Second, we provide a systematic evaluation of subspace and graph-embedded subspace-based variants of SVDD, demonstrating the central role of representation learning in mitigating the curse of dimensionality and stabilizing detection performance across diverse disease states. Third, we introduce a fully data-driven hyperparameter estimation strategy that infers all model parameters directly from intrinsic properties of the healthy training data, eliminating reliance on disease labels or exhaustive grid search in severely imbalanced data settings. Together, these contributions establish subspace learning based OCC as a robust, biologically grounded, and disease-agnostic framework for establishing a computational platform for high-dimensional plasma proteomics that can be used for discovering (the existence of) and detecting (the presence of) anomalies for future use in cohort screening or personalized medicine.

## 2 Results

We evaluated the proposed healthy-trained one-class learning framework on public plasma proteomics data generated using Olink’s PEA assays. The study included three independent cohorts: a healthy reference cohort, from which 70% of samples were used for model training and the remaining 30% were reserved for evaluation, multiple cancer cohorts and an independent COVID-19 cohort used exclusively for testing. The details of the datasets are provided in Section 4.

### One-class learning framework with data-driven hyperparameter estimation

An overview of the proposed healthy-profile-based one-class learning framework is presented in Fig. 1. Plasma samples are first profiled using Olink’s PEA technology, producing high-dimensional Normalized Protein Expression (NPX) measurement values for each individual. The proposed approach models the distribution of physiologically normal proteomic variation using OCC, where only healthy reference samples are used during training. To capture the dominant structure of healthy proteomic variation while reducing dimensionality, we employ subspace learning variants of SVDD. These methods jointly learn a projection from the original high-dimensional proteomic feature space into a lower-dimensional representation (the subspace) and construct a compact hyperspherical boundary that encloses the majority of healthy samples in this learned subspace. In addition to the standard SSVDD formulation, we also consider graph-embedded extensions that incorporate relationships between samples in order to preserve local geometric structure in the learned representation.

**Fig. 1:**
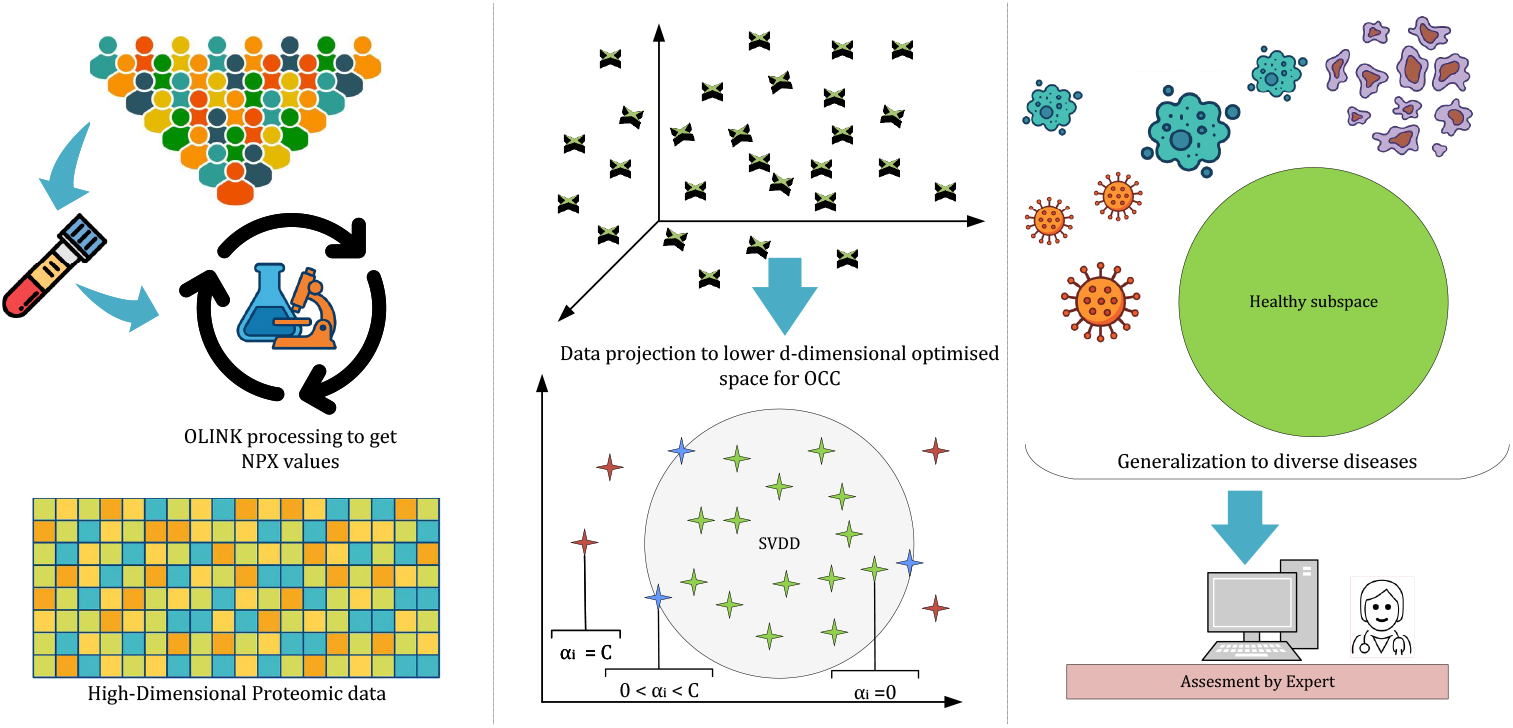
Conceptual overview of the healthy-profile-based subspace one-class learning framework. Plasma samples analyzed using Olink’s PEA technology to obtain high-dimensional protein abundance (NPX) values. A low-dimensional subspace is learned exclusively from healthy reference samples using subspace or graph-embedded SSVDD. A compact hyperspherical boundary is then constructed within the learned subspace to characterize normal proteomic variation. New samples from heterogeneous disease cohorts are projected into the learned representation space and classified as either consistent with the healthy manifold or anomalous. The framework enables disease-agnostic detection without using samples with disease labels during training.

Once the healthy reference model is established, new samples from independent disease cohorts are projected into the learned representation space, where samples located inside the learned boundary are considered consistent with the healthy proteomic manifold and those outside are flagged as anomalous. Because the model is trained exclusively on healthy data, the framework enables disease-agnostic detection of abnormal proteomic profiles without requiring disease-specific labels. To systematically evaluate how different subspace learning strategies influence this healthy-profile modeling, we consider several families of subspace-based OCC methods, including SSVDD and Newton-based SSVDD (NSSVDD) [23], which differ in the optimization strategy used to learn the projection matrix, as well as Graph-Embedded SSVDD (GESSVDD) [20] variants that incorporate structural relationships between samples through graph-based regularization. For the GESSVDD models, three graph constructions are considered: identity graphs (I), principal component-based graphs (PCA), and *k*-nearest neighbor graphs (kNN), each solved using Gradient-based (Gr), eigen-value decomposition based (Ei), and spectral regression based (Sr) solutions.

A critical design principle of the framework is that all model parameters are inferred exclusively from intrinsic properties of the healthy training data, without reliance on disease labels, cross-validation on diseased samples, or extensive grid search. This ensures that detection performance measured on cancer and COVID-19 cohorts reflects genuine deviations from the healthy proteomic manifold rather than implicit adaptation to disease-specific patterns. For the non-linear formulation, the kernel bandwidth parameter *σ* was inferred automatically using the median of pairwise distances among healthy samples, adapting naturally to the scale and dispersion of the healthy proteomic data and avoiding instability associated with manual kernel tuning in high-dimensional and imbalanced settings.

The dimensionality *d* of the learned subspace was determined directly from the variance structure of the healthy data. Singular Value Decomposition (SVD) was applied to the healthy reference cohort, and the smallest number of components explaining at least 95 percent of the total variance was retained, allowing effective model complexity to adapt to the intrinsic dimensionality of the healthy proteomic manifold. The regularization parameter (*β*) controlling boundary flexibility and subspace constraints was similarly derived from data-driven rules based on sample size and feature-wise variance. Graph-embedded variants additionally used neighborhood sizes that scaled logarithmically with the number of healthy samples, ensuring robust graph construction across cohort sizes. Together, this parameterization strategy eliminated the need for manual tuning, prevented information leakage from disease cohorts, and enabled fair and consistent comparison of subspace-learning based one-class models under realistic clinical constraints. The mathematical details of hyperparameter estimation are provided in Supplementary 1.

### Influence of model configuration on detection stability

The comparison of representative configurations from each methodological family further illustrates the influence of model configuration on detection stability (Fig. 2). In the healthy versus COVID-19 evaluation setting, most configurations achieved high GM values above 0.92, indicating consistent discrimination between healthy samples and COVID-19 proteomic profiles despite the absence of COVID-19 profiles during training. In particular, graph-embedded configurations based on *k*-nearest neighbor structure (GESSVDD-kNN-Ei-min) achieved the highest performance, reaching GM values close to 0.95 under both linear and non-linear formulations. Standard subspace approaches (SSVDD and NSSVDD) also showed stable behavior, although with slightly lower GM values.

**Fig. 2:**
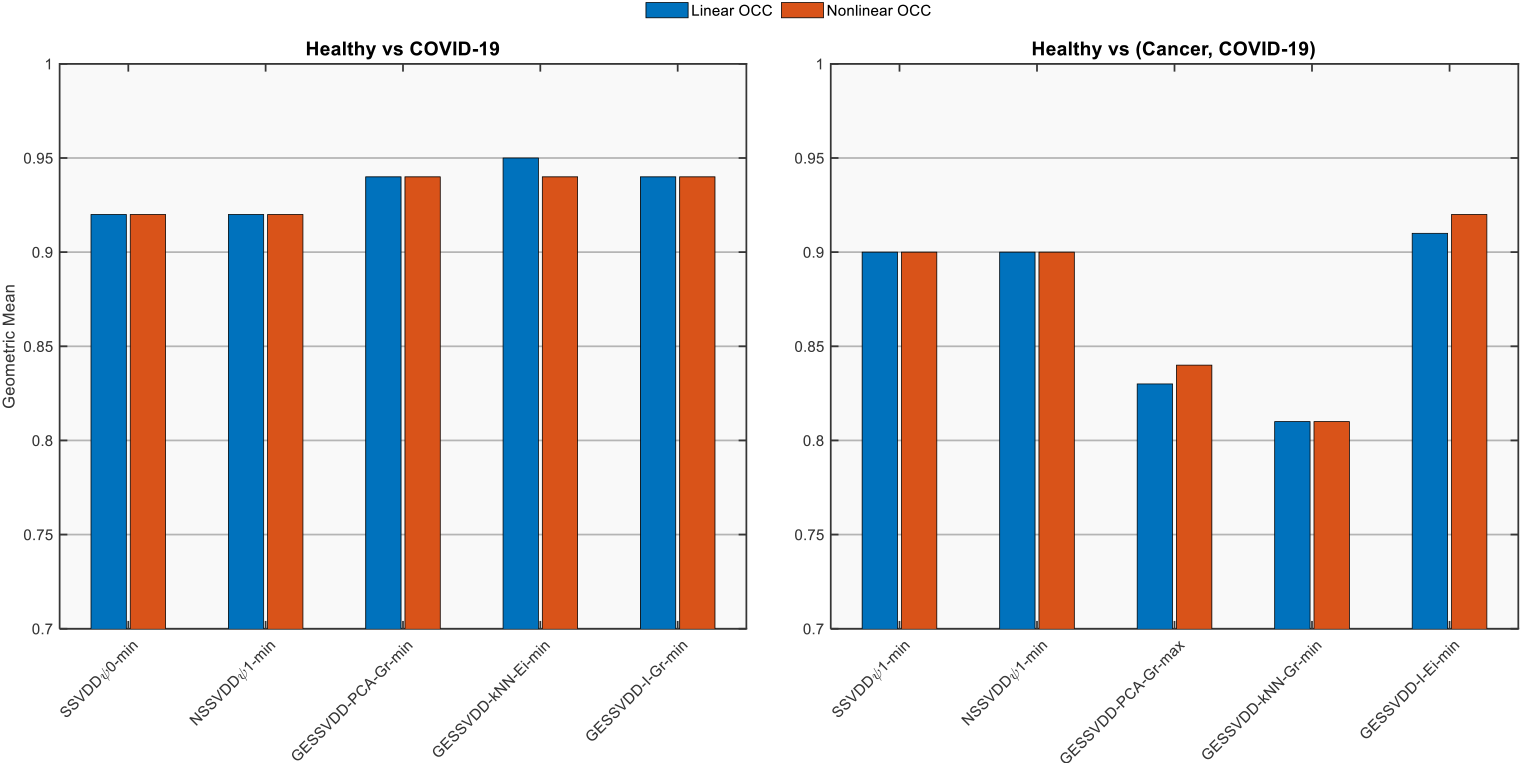
Comparison of healthy-trained one-class models across disease settings. Geometric Mean (GM) performance of representative OCC methods trained exclusively on healthy plasma proteomic profiles and evaluated on independent disease cohorts. The left panel shows performance for discrimination between healthy and COVID-19 samples, while the right panel shows performance for discrimination between healthy and combined cancer and COVID-19 samples. For each methodological family (SSVDD, NSSVDD, and graph-embedded SVDD variants), the best-performing configuration is shown under linear and non-linear formulations. All models use identical training data and data-driven parameter estimation, enabling direct comparison of detection behavior across disease contexts.

By contrast, greater variability between configurations emerged in the more challenging healthy versus pooled cancer and COVID-19 evaluation setting. While baseline subspace models (SSVDD and NSSVDD) maintained stable GM values around 0.90, several graph-embedded variants showed reduced performance depending on the graph construction. For example, PCA-based and kNN graph formulations exhibited noticeable drops in GM, reflecting sensitivity to structural constraints imposed during representation learning. Among the evaluated configurations, the identity-graph formulation (GESSVDD-I-Ei-min) achieved the highest detection performance in the pooled disease setting, reaching GM values above 0.91.

Across both evaluation settings, differences between linear and non-linear formulations were relatively small, indicating that the primary determinants of detection stability arise from the structural configuration of the learned subspace rather than the introduction of non-linear transformations. These observations support the conclusion that regularization strategy and graph construction play a central role in shaping the stability of healthy-trained one-class models in high-dimensional plasma proteomic data. We provide more detailed results tables along with other evaluated metrics in Supplementary 2: Detailed Performance Results for Disease Detection Tasks.

### Healthy-trained one-class models detect disease-associated proteomic deviations

To characterize the detection behavior of the selected models, we examined two complementary evaluation settings that assess the robustness of the learned healthy representation under different disease contexts. The first setting evaluates discrimination between healthy samples and an independent COVID-19 cohort, providing a controlled assessment of proteomic deviations in infectious diseases. The second setting pools cancer and COVID-19 samples into a single heterogeneous disease group, creating a more challenging scenario in which multiple disease processes perturb the plasma proteome. The graph embedded model GESSVDD-kNN-Ei-min showed the strongest performance in the healthy versus COVID-19 setting, whereas GESSVDD-I-Ei-min exhibited the most stable performance when cancer and COVID-19 samples were evaluated jointly. Figure 3 summarizes the full set of performance metrics for these representative models under both linear and non-linear formulations.

**Fig. 3:**
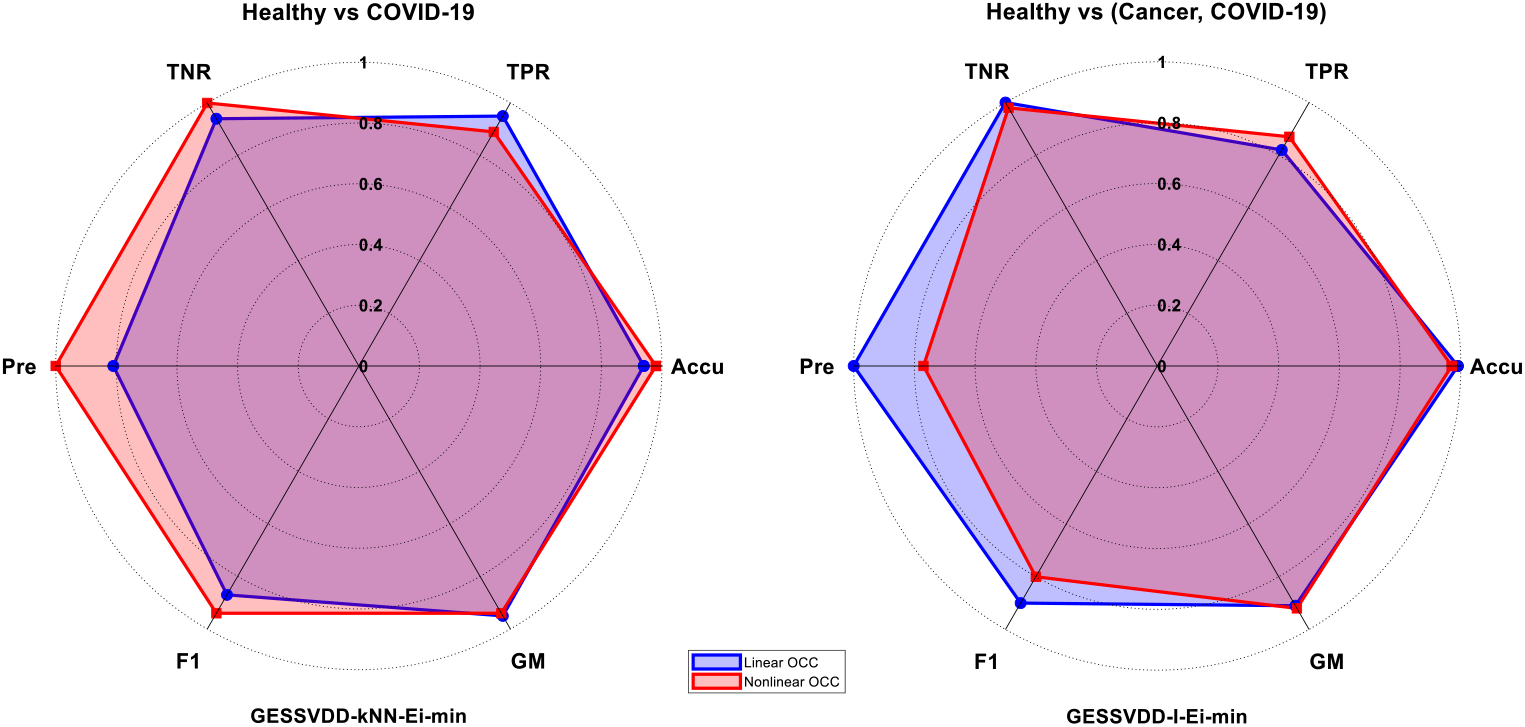
Radar chart comparison of representative healthy-trained one-class models across disease evaluation settings. Each radar plot summarizes six performance metrics (Accuracy, True Positive Rate, True Negative Rate, Precision, F1-score, and GM) for linear and non-linear formulations of selected subspace one-class models. The left panel corresponds to the healthy versus COVID-19 evaluation setting using the GESSVDD-kNN-Ei-min configuration, while the right panel corresponds to the pooled healthy versus cancer and COVID-19 evaluation setting using the GESSVDD-I-Ei-min configuration. All models were trained exclusively on healthy plasma proteomic profiles and evaluated on independent disease cohorts using identical training partitions and fully data-driven parameter estimation. Shaded regions represent linear and non-linear OCC variants, highlighting differences in detection behavior across disease contexts.

In the healthy versus COVID-19 evaluation setting, the locally structured GESSVDD-kNN-Ei-min model achieved consistently strong performance across all metrics. As shown in Fig. 3 (left), both linear and non-linear formulations maintained high accuracy and GM values while achieving near perfect true positive rates, indicating that healthy reference samples were reliably identified as healthy. True negative rates were also high, demonstrating that COVID-19 samples were correctly identified as disease relative to the healthy reference cohort. The non-linear formulation provided modest improvements in precision and F1 score, indicating a trade-off between balanced detection performance and class-specific discrimination when nonlinear transformations are incorporated.

The second radar plot in Fig. 3 summarizes performance in the pooled healthy versus cancer and COVID-19 evaluation setting using the GESSVDD-I-Ei-min configuration. In this heterogeneous disease scenario, both linear and non-linear formulations maintained high accuracy and true negative rates, indicating reliable identification of disease samples across diverse oncological and infectious conditions. The true positive rate was moderately reduced compared to the COVID-only evaluation, reflecting the increased difficulty of preserving the healthy boundary when multiple disease cohorts are considered simultaneously. Despite this added heterogeneity, the model maintained balanced performance across metrics, achieving GM values close to 0.9.

Within the COVID-19 cohort, we further examined a subgroup of clinically “healthy-like” individuals, comprising eleven samples, to assess how the learned healthy manifold behaves under moderate distributional shift. In the COVID-only evaluation setting, most baseline one class configurations classified these eleven samples as disease relative to the healthy reference distribution, with none of the individuals correctly retained as healthy in most configurations. In contrast, the locally constrained GESSVDD-kNN-Ei-min model retained up to 7 of the 11 healthy-like COVID-19 samples within the learned healthy boundary. When cancer and COVID-19 samples were pooled, the evaluation was performed using the 62 protein assays shared across all cohorts. Under this reduced cross-cohort feature space, none of the evaluated models retained the healthy-like COVID-19 individuals. Because the pooled analysis relies only on assays consistently measured across datasets, the reduced feature set may limit the representation of subtle proteomic patterns that distinguish these individuals from the disease cohorts. As a result, these samples were systematically classified as disease relative to the healthy reference manifold, suggesting that informative signals required to retain them as healthy may be present in assays not shared across cohorts.

Across both evaluation settings, true negative rates remained consistently high, indicating that the learned hyperspherical boundary effectively distinguishes disease-associated proteomic profiles from the healthy reference distribution. This consistent performance across heterogeneous cohorts demonstrates that the learned representation captures proteomic deviations associated with both infectious and oncological conditions. True positive rates also remained high, confirming that the structure of the healthy reference manifold is largely preserved. Differences in overall detection performance were therefore mainly reflected in small variations in the balance between these metrics across model configurations. Robustness to such dataset shifts is critical for clinical deployment, where differences between training and deployment populations are unavoidable [24].

### Proteomic anomaly scores reflect clinical severity in the COVID-19 cohort

To examine whether the proteomic anomaly scores produced by the healthy-trained one-class model captured graded clinical severity, we stratified Day 0 SVDD distances by acuity score within the COVID-19 cohort. Acuity scores reflect the highest severity recorded within the Day 0 window (enrollment plus 24 hours), ranging from 1 (death) to 5 (discharged or not hospitalised). Median distances to the healthy 9 proteomic manifold centre decreased monotonically from Acuity 2 (intubated or ventilated) to Acuity 5 (discharged), consistent with greater proteomic deviation from the healthy reference distribution in more severely ill patients. Notably, this severity gradient emerged without any disease severity information being available during model training, demonstrating that the learned healthy subspace captures biologically meaningful structure that generalises to graded pathological states. Healthy-like individuals, defined by the absence of recorded comorbidities, showed distances concentrated near or below the decision boundary, with those retained within the healthy manifold pre-dominantly found at lower acuity levels. These results further support the conclusion that the GESSVDD-kNN model detects clinically relevant proteomic deviations in a disease-agnostic manner (Fig. 4).

**Fig. 4:**
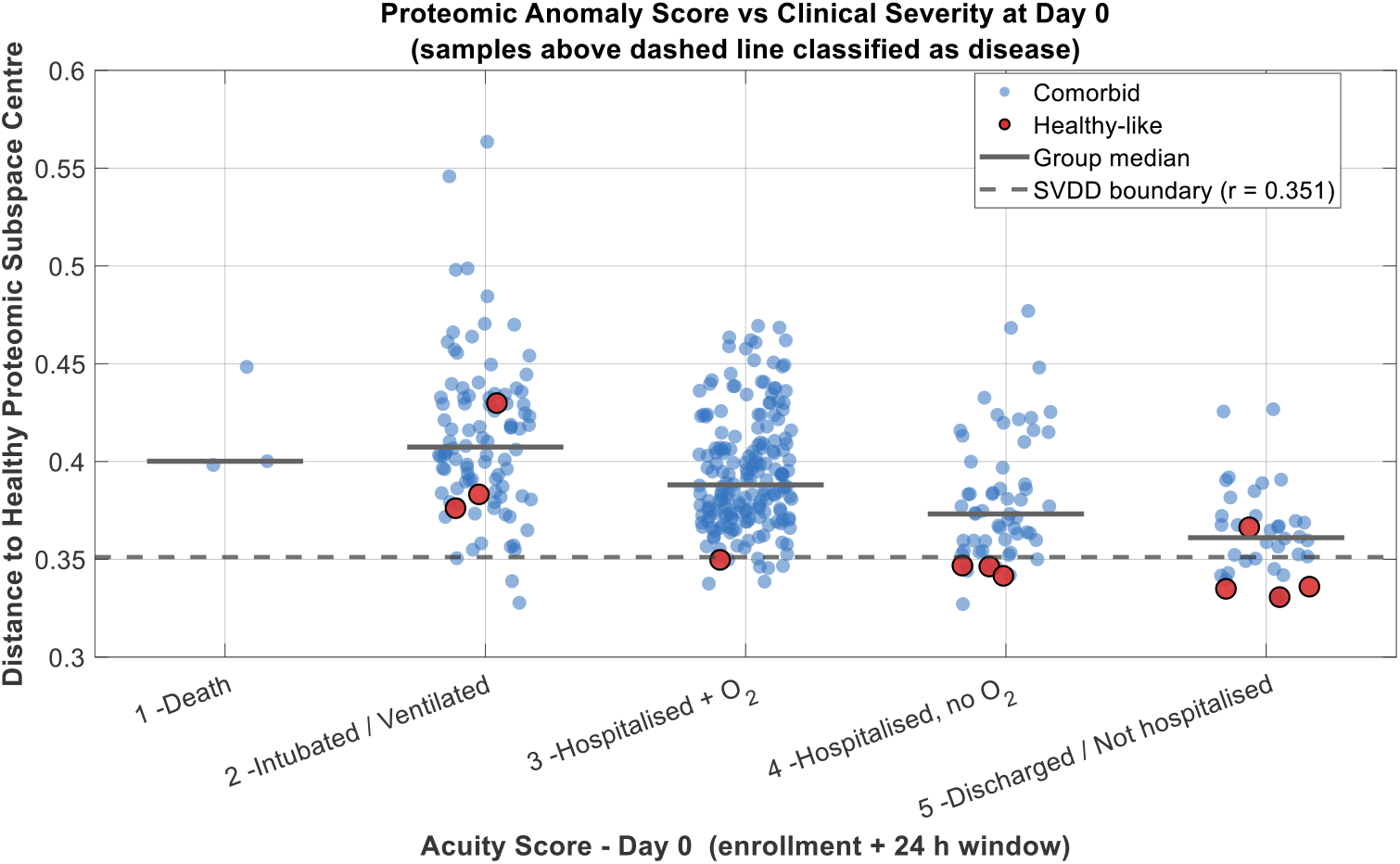
Proteomic anomaly scores stratified by Day 0 clinical acuity in the COVID-19 cohort. Each point represents an individual plasma proteomics sample projected into the learned subspace of GESSVDD-kNN and scored by distance to the healthy proteomic manifold centre. Acuity scores reflect the highest severity recorded within the Day 0 window (enrollment plus 24 hours): 1 = death, 2 = intubated or ventilated survived, 3 = hospitalised with supplementary oxygen survived, 4 = hospitalised without supplementary oxygen survived, 5 = discharged or not hospitalised survived. Comorbid individuals (blue) and healthy-like individuals (red), defined by the absence of recorded comorbidities, are shown separately. The dashed line indicates the SVDD decision boundary (r = 0.351); samples above this threshold are classified as disease relative to the healthy reference manifold. Horizontal bars indicate group medians per acuity category. All models were trained exclusively on healthy reference plasma proteomic profiles, with no disease severity information used during training or parameter estimation.

### Assessment of demographic structure in healthy proteomic profiles

To assess whether healthy-trained one-class models inadvertently encode demographic structure unrelated to disease, we performed auxiliary experiments within the healthy reference cohort using available metadata for sex and ethnicity. One-class models were trained with each demographic group alternately treated as the target (“healthy”) class to evaluate whether healthy plasma proteomic profiles exhibit separable structure along these axes.

Across all evaluated subspace and graph-embedded formulations, detection performance for both sex and ethnicity remained near chance level and exhibited high instability across configurations (Fig. 5), collectively indicating the absence of consistent separable demographic structure in the learned healthy subspaces. Rather than encoding sex-or ethnicity-related variation, the learned representations appear to capture proteomic signals that are largely orthogonal to these demographic axes, suggesting that detected deviations from the healthy manifold reflect biological perturbation rather than sociodemographic differences. Complete performance metrics for all configurations and demographic groupings are reported in Supplementary 3.

**Fig. 5:**
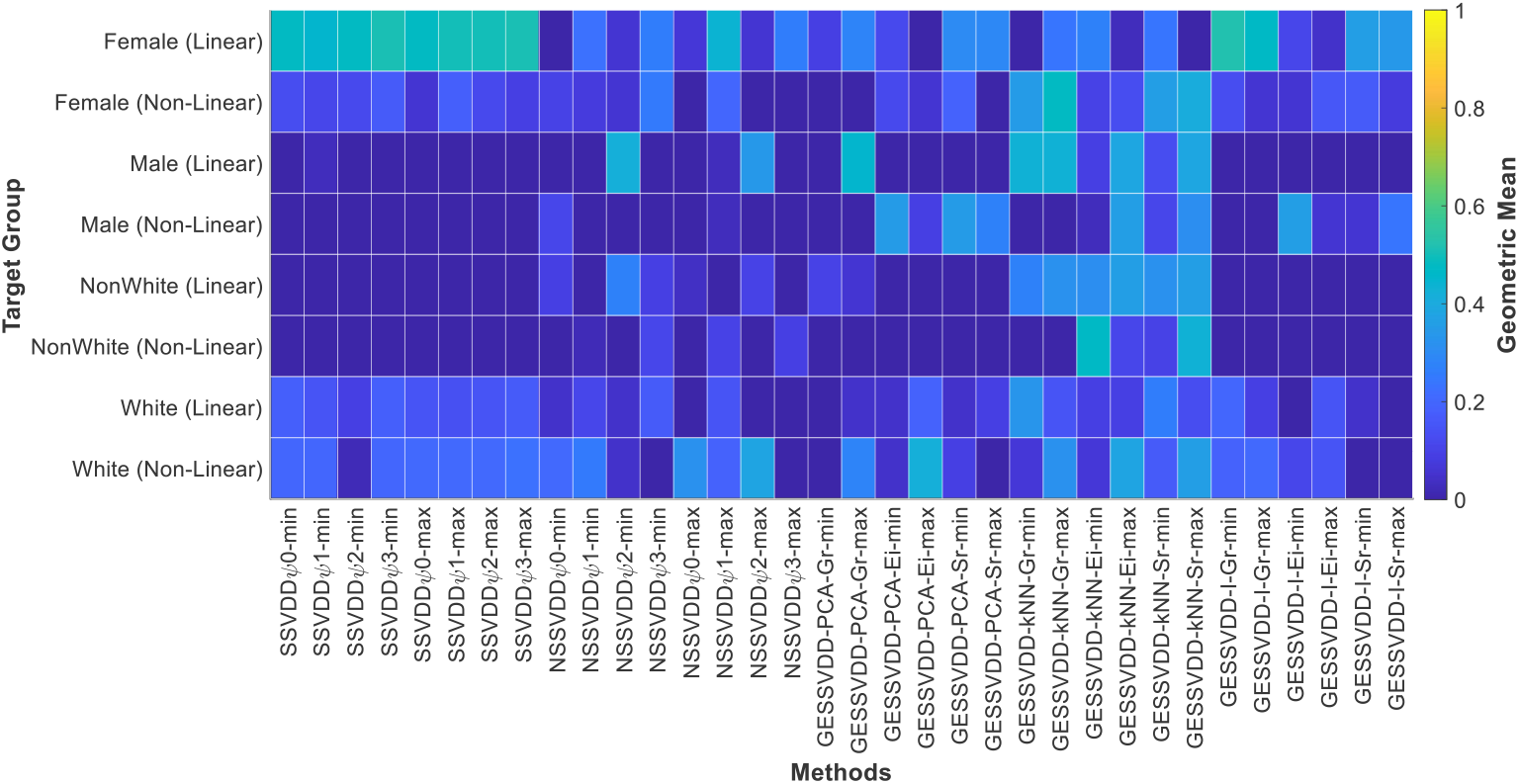
Heatmap of GM performance across demographic target classification tasks. Rows correspond to target groups and model type (Linear and Non-Linear OCC), while columns represent the evaluated methods. Color intensity reflects the magnitude of the GM, highlighting performance variability across gender and ethnicity targets.

## 3 Discussion

This study investigates one-class modeling of healthy plasma proteomic profiles in high-dimensional proteomic space as a general strategy to detect any disease that manifests as a departure from the healthy profile in an a priori unknown direction. By training all models solely on a healthy reference cohort and evaluating performance across multiple cancer types and an independent COVID-19 cohort, we present a systematic assessment of how subspace one-class learning generalizes from normal proteomic variation to diverse pathological states. The results demonstrate that modeling the healthy proteomic manifold in a learned low-dimensional space is sufficient to capture deviations associated with both cancer and viral infection, supporting the practical utility of healthy-trained one-class approaches in settings where diseased samples are scarce, heterogeneous, or too novel to permit incorporation of pathological patterns during training.

A central finding of this work is the critical role of representation learning in stabilizing one-class detection performance in plasma proteomics. All evaluated models operate in learned subspaces rather than the original high-dimensional feature space, enabling suppression of noise and redundancy while preserving coordinated biological variation. Across disease contexts, subspace-based formulations yielded compact and structured representations of healthy proteomic variation, resulting in more balanced detection behavior and reduced sensitivity to spurious fluctuations. This behavior is biologically plausible, as coordinated protein responses are driven by regulatory and signaling processes and are therefore expected to reside on low-dimensional manifolds embedded within the high-dimensional measurement space. Learning such representations allows one-class models to more reliably expose meaningful deviations from the biological norm in the presence of technical or stochastic variation.

Evaluation on an independent COVID-19 cohort highlights the disease-agnostic nature of healthy-trained subspace modeling. Despite the absence of any pathological samples during training or parameter estimation, the learned healthy subspaces supported consistent detection of proteomic deviations associated with acute viral infection. This generalization across fundamentally different disease mechanisms indicates that subspace-based models capture intrinsic properties of physiological plasma proteome organization rather than disease-specific signatures. Graph-embedded variants demonstrated particular robustness under this distributional shift, suggesting that explicitly encoding geometric structure within the healthy reference manifold improves stability when encountering previously unseen pathological states. Complementing these findings, stratification of anomaly scores by Day 0 clinical acuity revealed a monotonic decrease in median distance to the healthy manifold centre from the most severe patients (intubated or ventilated) to those discharged or not hospitalised. This severity gradient emerged without any clinical severity information being used during training, indicating that the learned healthy subspace captures biologically meaningful variation that extends beyond binary disease detection to reflect graded pathological perturbation of the plasma proteome.

At the same time, no single subspace-based modeling strategy uniformly dominated across all disease cohorts, graph constructions, or regularization settings. Instead, different methods exhibited characteristic trade-offs between sensitivity and specificity, reflecting their underlying assumptions about data geometry and variance structure. This observation underscores the importance of comparative evaluation within a unified framework rather than reliance on a single algorithmic choice. In practice, method selection should be informed by the intended application, acceptable false positive rates, and the expected diversity of disease-associated proteomic perturbations.

Several limitations of this study warrant consideration. The evaluation is limited to plasma proteomics data generated using a single platform; therefore, the extent to which these findings generalize to other proteomic technologies or tissue types remains to be established. Moreover, while healthy-only training avoids biases introduced by incomplete or heterogeneous disease labels, it does not capture disease-specific heterogeneity, progression dynamics, or temporal effects, which may be important for prognosis or subtype differentiation. Future work could extend this framework to longitudinal designs, multi-omics integration, and semi-supervised or adaptive extensions that incorporate limited disease information without compromising generalization. An important practical consideration for population-scale screening models is the risk of unintended demographic stratification. This concern mirrors broader findings in clinical machine learning, where demographic bias can arise even in the absence of explicit labels [25, 26]. Using available metadata from the healthy reference cohort, we evaluated whether one-class models trained on healthy plasma proteomics implicitly separate individuals by sex or ethnicity. The consistently poor and unstable performance observed across all formulations indicates that healthy proteomic variation, as captured by the learned subspaces, does not strongly encode demographic structure. This behavior is desirable in a disease-agnostic screening context, as it reduces the risk of demographic bias and supports the interpretation that detected deviations reflect fundamental pathological rather than sociodemographic variation.

In summary, this work demonstrates that one-class subspace modeling of healthy plasma proteomic profiles provides a robust and biologically grounded approach for disease detection in high-dimensional proteomics. By systematically evaluating subspace and graph-embedded variants across cancer and COVID-19 cohorts using fully data-driven parameterization, we demonstrated the strengths and potential utility of representation learning within the one-class framework. The observed correspondence between proteomic anomaly scores and clinical severity further supports the translational relevance of this approach. These findings motivate continued development of interpretable and scalable subspace one-class methods for omics profiling in medicine, with potential applications in healthy-reference modeling, disease-agnostic population screening, and early detection of clinically silent pathological states. More broadly, recent work in anomaly detection has emphasized the suitability of one-class learning paradigms for biomedical applications in which abnormal states are heterogeneous, evolving, and incompletely characterized [27].

## 4 Methods

### Study design and one-class learning framework

This study adopts the OCC framework in which models are trained exclusively on proteomic profiles from clinically healthy individuals and evaluated on independent disease cohorts. The objective is to learn a compact representation of normal proteomic variation and identify deviations associated with pathological states without using disease-specific information during training. This design enables an unbiased assessment of how different one-class modeling strategies generalize from healthy reference data to heterogeneous disease conditions. All experiments were conducted using plasma proteomics data generated with Olink’s PEA, reported as NPX values. Cancer and COVID-19 cohorts were used exclusively for evaluation and were never included in model training or parameter estimation.

### Subspace Support Vector Data Description

All OCC models evaluated in this study operate in a learned low-dimensional subspace rather than the original high-dimensional proteomic feature space. The proposed framework, referred to as Parameter-Free Subspace Support Vector Data Description (PFSSVDD), infers all model parameters directly from the healthy training data without cross-validation or disease labels, and serves as the overarching methodology within which all evaluated subspace and graph-embedded variants are unified. Let **x**_*i*_ ∈ℝ^*D*^ denote standardized NPX feature vectors from the healthy training cohort. A projection matrix **Q** ∈ℝ^*d×D*^ with *d* ≤*D* maps samples into a lower-dimensional representation

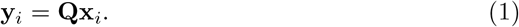

Within this subspace, a hyperspherical boundary is constructed to characterize normal proteomic variation. The SSVDD formulation jointly learns the projection matrix and the hypersphere parameters by minimizing

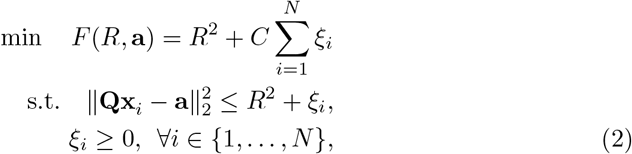

where *R* denotes the radius of the hypersphere and **a** ∈ℝ^*d*^ represents its center in the projected space. The parameter *C >* 0 controls the trade-off between minimizing the hypersphere volume and penalizing samples outside the boundary. Slack variables *ξ*_*i*_ allow certain training samples to lie outside the hypersphere, improving robustness to potential outliers. The augmented Lagrangian formulation of the objective is

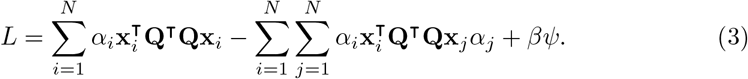

Optimising (3) yields Lagrange multipliers *α*_*i*_ associated with each training sample. Data points satisfying 0 *< α*_*i*_ *< C* define the hypersphere boundary and are referred to as support vectors. The regularization functional *ψ* characterizes the variance structure of healthy proteomic profiles within the learned *d*-dimensional subspace and *β* controls the importance of the regularization term. The regularization term *ψ* is defined as

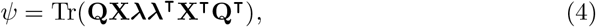

where Tr(·) denotes the trace operator and ***λ*** ∈ℝ^*N*^ is a selection vector controlling which samples contribute to the estimation of variance in the projected subspace. By modifying ***λ***, different variants of the SSVDD formulation can be obtained, each imposing a different geometric constraint on the representation of healthy proteomic variation. The following different variants are used in this study

- *ψ*_0_: No explicit regularization term is used, and the projection matrix is learned solely from the hypersphere data description objective.
- *ψ*_1_: All training samples contribute to the variance estimation, encouraging the learned subspace to preserve the overall distribution of healthy proteomic profiles.
- *ψ*_2_: Samples located on or outside the hypersphere boundary are used in the regularization term, emphasizing the representation of boundary points that define the limits of the healthy data description.
- *ψ*_3_: Only support vectors (0 *< α*_*i*_ *< C*), which lie on the hypersphere boundary, contribute to the regularization term.

These regularization variants allow different assumptions about the structure of healthy proteomic variation to be incorporated into the learned subspace, enabling systematic evaluation of how variance preservation versus boundary-focused representations influence anomaly detection performance [19, 28]. The update of the projection matrix **Q** is obtained from the gradient of (3), defined as:

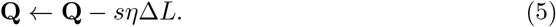

Here, *η* denotes the learning rate parameter and Δ*L* represents the gradient with respect to the projection matrix **Q**. The variable *s* ∈{™1, 1} determines the direction of the update. Specifically, *s* = 1 corresponds to a maximization strategy, whereas *s* = ™1 corresponds to a minimization strategy. In addition to the gradient-based optimization used in SSVDD, we also consider a Newton-based optimization strategy for learning the projection matrix. While gradient descent relies only on first-order information, Newton’s method incorporates second-order curvature information through the Hessian of the objective function, which can improve convergence stability and optimization efficiency.

### Graph-Embedded Subspace Support Vector Data Description

GESSVDD extends the SSVDD formulation by incorporating structural relationships among samples during subspace learning. Let **L**_*x*_ denote the graph Laplacian constructed from the healthy cohort. The projected scatter matrix is defined as

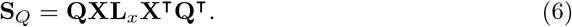

The projection matrix defining the subspace is given by 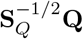, which preserves structural relationships encoded in the graph while learning the low-dimensional representation. Three graph constructions were evaluated:

- Identity graph (I), imposing no additional structural constraints,
- Principal component-based graph (PCA), emphasizing global variance structure,
- *k*-nearest neighbor graph (kNN), preserving local manifold geometry.

For each graph construction, three solution methods were considered: a graph-regularized formulation (Gr), an eigen-decomposition-based solution (Ei), and a spectral regression formulation (Sr). Model names reported in the Results section reflect the graph type and the corresponding solution method. All models were evaluated under both linear and nonlinear formulations. Linear variants operate directly on standardized NPX features, whereas nonlinear variants employ an RBF kernel mapping implemented through the Nonlinear Projection Trick (NPT) [29]. This approach allows subspace learning in an induced feature space without explicit kernel matrix optimization while preserving the SVDD hyperspherical data description framework. Together, these formulations define a unified family of subspace-based one-class models differing in optimization strategy, regularization scheme, and structural constraints on the learned representation.

### Datasets and preprocessing

Plasma proteomics data used in this study comprised three independent cohorts: a healthy reference cohort, multiple cancer cohorts, and an independent COVID-19 cohort (Supplementary Fig. S1). All proteomic measurements were generated using Olink’s PEA and reported as NPX values. The study followed a strict OCC design, in which only healthy samples were used for model training and hyperparameter estimation. All cancer and COVID-19 samples were withheld from training and used exclusively for evaluation.

The healthy reference cohort consisted of plasma samples from clinically healthy individuals without known cancer, infectious disease, or major chronic conditions. This cohort served as the sole source of training data for all models. Healthy samples were used to estimate feature-wise normalization parameters, determine subspace dimensionality, construct graph embeddings, and infer all model hyperparameters using fully data-driven procedures without access to disease labels.

Cancer data comprised multiple oncological cohorts spanning diverse cancer types [30]. For evaluation, cancer samples were pooled into a single disease group to assess overall detection performance, while individual cancer types were retained for descriptive analyses and stratified reporting. No cancer samples were used during model fitting, normalization, or parameter estimation.

An independent COVID-19 cohort [31] was included to evaluate cross-disease generalization to an unseen infectious disease. Clinical metadata were used to stratify individuals within the COVID-19 cohort into COVID-19 healthy-like and COVID-19 comorbid groups. A composite clinical indicator was constructed as the logical OR across recorded diagnoses of COVID-19 infection, cardiovascular disease, lung disease, kidney disease, diabetes, hypertension, and immunological disorders. Individuals with no recorded conditions were classified as COVID-19 healthy-like, whereas individuals with at least one recorded condition were classified as COVID-19 with comorbidities. This stratification was used exclusively for post hoc analysis and visualization and did not influence model training, normalization, or decision boundary construction. A two-dimensional PCA projection of all cohorts onto the first two principal components, computed from the 62 shared protein assays, is provided in Supplementary Fig. S2.

To ensure consistent input dimensionality across cohorts, only protein assays shared across the healthy, cancer, and COVID-19 datasets were retained. Olink feature identifiers were mapped to assay names prior to alignment. Protein features containing missing values in any cohort were removed globally to enforce a common feature space and to avoid cohort-specific bias during training and evaluation. For the healthy versus COVID-19 evaluation, this procedure yielded 1,415 shared protein assays, whereas the pooled evaluation including all three cohorts retained 62 shared assays, reflecting the more stringent overlap requirement when cancer cohort data were incorporated.

Feature normalization was performed using mean and standard deviation estimates computed exclusively from the healthy training subset. These normalization parameters were applied unchanged to all evaluation samples, including held-out healthy samples, cancer cohorts, and the COVID-19 cohort, thereby preventing information leakage from disease data while ensuring comparable feature scaling across cohorts.

### Evaluation protocol

Healthy reference samples were randomly partitioned using a stratified holdout strategy, with 70% of healthy samples used for model training and 30% reserved as an internal healthy test set. The training subset was used exclusively for model fitting, normalization parameter estimation, subspace learning, graph construction, and hyperparameter inference.

For evaluation, the held-out 30% healthy samples were combined with all cancer samples and all COVID-19 samples to form the test set. This design enabled simultaneous assessment of false positive behavior on unseen healthy individuals and detection performance on heterogeneous disease cohorts under a unified and unbiased evaluation protocol. The data partitioning procedure was repeated multiple times to assess the stability of model performance.

Detection performance was quantified using accuracy, sensitivity, specificity, precision, F1-score, and GM. GM was used as the primary performance metric due to its balanced treatment of detection performance across classes in the presence of class imbalance. Performance metrics were averaged across repeated splits to ensure robustness.

## Data Availability

Cancer cohort data are publicly available from the EBI BioStudies repository under accession S-BSST935 (https://www.ebi.ac.uk/biostudies/studies/S-BSST935). COVID-19 cohort data were generated as part of the Massachusetts General Hospital (MGH) COVID-19 study. Access to the NPX-level proteomic data and associated clinical metadata is subject to controlled access and can be requested through the study information page provided by Olink (https://insight.olink.com/olink-data/covid), subject to approval by the data custodians and applicable data use agreements. The healthy plasma proteomics reference cohort used for training the one-class models was generated using Olink PEA technology. Due to ethical approvals, participant consent restrictions, and data use agreements, the raw NPX-level data for the healthy reference cohort are not publicly available. Access may be granted upon reasonable request to the corresponding author and is subject to approval by the relevant data custodians and institutional review boards.

## Code availability

The source code supporting the methods and experiments presented in this study is publicly^1^ available on GitHub at https://github.com/fahadsohrab/pfssvdd/. The repository includes implementations of the subspace and graph-embedded one-class models, along with scripts for preprocessing, evaluation, and reproduction of the reported results.

## Acknowledgements

The authors acknowledge support from institutional research funding and infrastructure provided by the University of Eastern Finland. The authors thank the data contributors and study participants whose publicly available and controlled-access proteomics datasets made this work possible. The use of Olink’s PEA proteomics data from the cancer and COVID-19 cohorts is gratefully acknowledged. This work was supported by the Jane and Aatos Erkko Foundation. The authors wish to acknowledge UEF Bioinformatics Center (Biocenter Finland) and CSC IT Center for Science, Finland, for computational resources.

## Author contributions

F.S. designed the methodology, performed the experiments, and wrote the manuscript.

A.K. and V.A. assisted with data analysis and computational implementation. A.M. provided the healthy reference proteomics dataset, associated technical expertise in Olink PEA data generation, and contributed to study design. V.H. contributed to methodological development, algorithmic design, and critical review of the manuscript.

M.H. and S.H. contributed equally as co-senior authors, providing joint scientific leadership, conceptual development of the study framework, and supervision of the research. All authors contributed to the interpretation of the results, critically reviewed the manuscript, and approved the final version.

## Supplementary 1: Hyperparameter Estimation

This supplementary provides the complete mathematical specification of the fully data-driven hyperparameter estimation strategy used in all one-class models. All parameters are inferred exclusively from the healthy training data without cross-validation, disease labels, or manual tuning. This design ensures strict separation between training and evaluation cohorts and prevents information leakage from disease samples. The resulting parameterization framework enables reproducible and unbiased comparison across linear, non-linear, and graph-embedded subspace formulations. Let **X** = [**x**_1_, …, **x**_*N*_ ] ∈ℝ^*D×N*^ denote the standardized training data matrix containing only samples from the target class.

### Kernel bandwidth and nonlinear projection

When a nonlinear mapping is employed, the proposed framework uses the Nonlinear Projection Trick (NPT). The kernel used in this work is RBF kernel as follows:

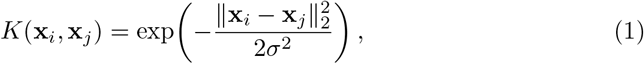

where *σ >* 0 denotes the kernel bandwidth parameter controlling the locality of the induced similarity measure.

Let

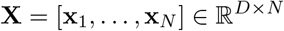

denote the standardized training data matrix consisting exclusively of target-class samples. The pairwise Euclidean distance between samples **x**_*i*_ and **x**_*j*_ is defined as

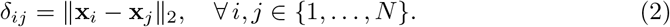

The kernel bandwidth is computed using the median distance heuristic:

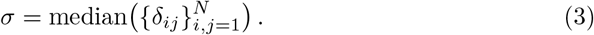

This choice provides a robust, data-adaptive estimate of the kernel scale, ensuring that the nonlinear mapping captures the typical geometric structure of the target-class data. The median heuristic avoids sensitivity to extreme distances and eliminates the need for cross-validation, which is particularly important in one-class learning scenarios. Within the NPT framework, the kernel-induced representation is computed explicitly, allowing subsequent subspace learning and SVDD optimization to be performed in the transformed space without requiring direct kernel matrix optimization. As a result, the nonlinear structure of the data is preserved while maintaining computational efficiency.

### Subspace dimensionality

The dimensionality of the learned subspace is selected using an energy preservation criterion based on singular value decomposition (SVD). Let

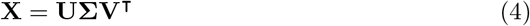

be the compact SVD of the training data. The subspace dimension *d* is chosen as

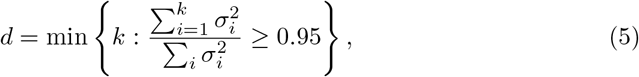

ensuring that at least 95% of the total variance is preserved.

### SVDD regularization parameter

The regularization parameter *C* of SVDD is derived from the *v*-SVDD formulation. Given a fixed outlier fraction *v* = 0.1, *C* is set as

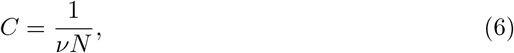

where *N* is the number of target-class training samples.

### Subspace regularization weight

For variants including an explicit subspace regularization term, the corresponding weight *β* is computed adaptively from the data. Let

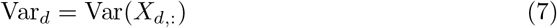

denote the variance of the *d*-th feature. The regularization weight is defined as

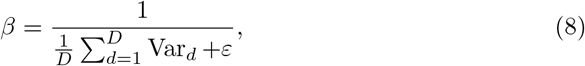

where *ε* = 10^*−*6^ is a small constant for numerical stability. This choice makes the regularization weight data-adaptive and scale-aware. When the average variance of the input features is large, the regularization weight *β* becomes smaller, preventing the regularization term from dominating the optimization objective. Conversely, for data with low variance, a larger *β* strengthens the regularization effect, promoting stable and well-conditioned subspace learning. Importantly, this formulation ensures that the magnitude of the regularization term is automatically aligned with the intrinsic variability of the data, eliminating the need for manual tuning or cross-validation. This is particularly desirable in one-class learning settings, where labeled validation data are typically unavailable or severely limited.

### Graph neighborhood size

For graph-based variants employing *k*-nearest neighbor graphs, the neighborhood size is determined automatically as

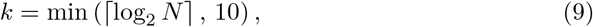

ensuring a balance between locality and robustness.

All hyperparameters are therefore estimated directly from the training data, eliminating the need for cross-validation and making the proposed method particularly suitable for one-class learning scenarios.

## Supplementary 2: Detailed Performance Results for Disease Detection Tasks

This supplementary reports the complete performance metrics for all evaluated one-class classification methods in the healthy versus COVID-19 setting (Table 1) and the pooled healthy versus cancer and COVID-19 setting (Table 2). Results are shown for both linear and non-linear formulations. Metrics include accuracy (Accu), true positive rate (TPR), true negative rate (TNR), precision (Pre), F1-score (F1), and geometric mean (GM).

**Table S1:**
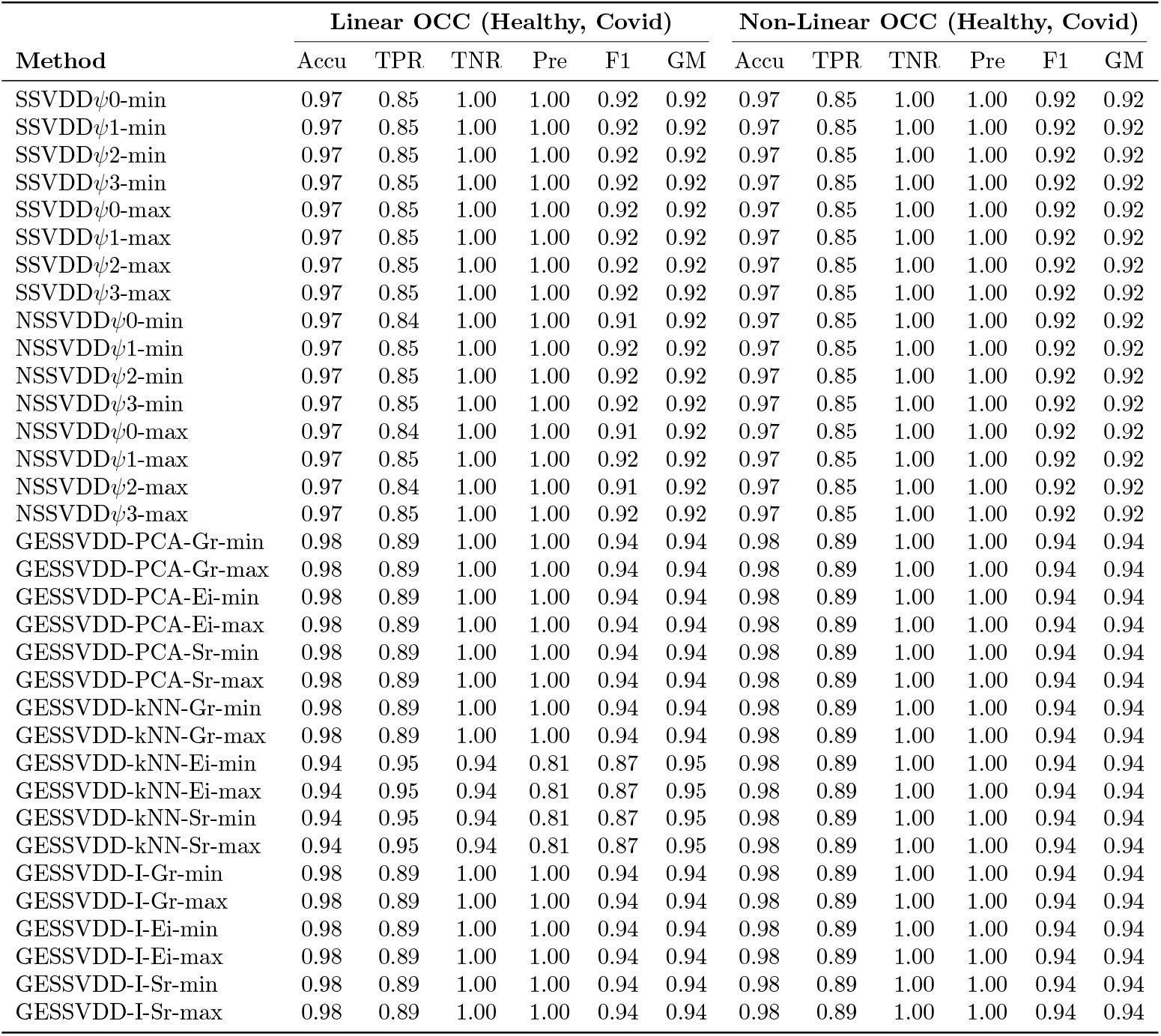
Performance comparison of Linear and Non-Linear OCC methods for (Healthy, Covid).

**Table S2:**
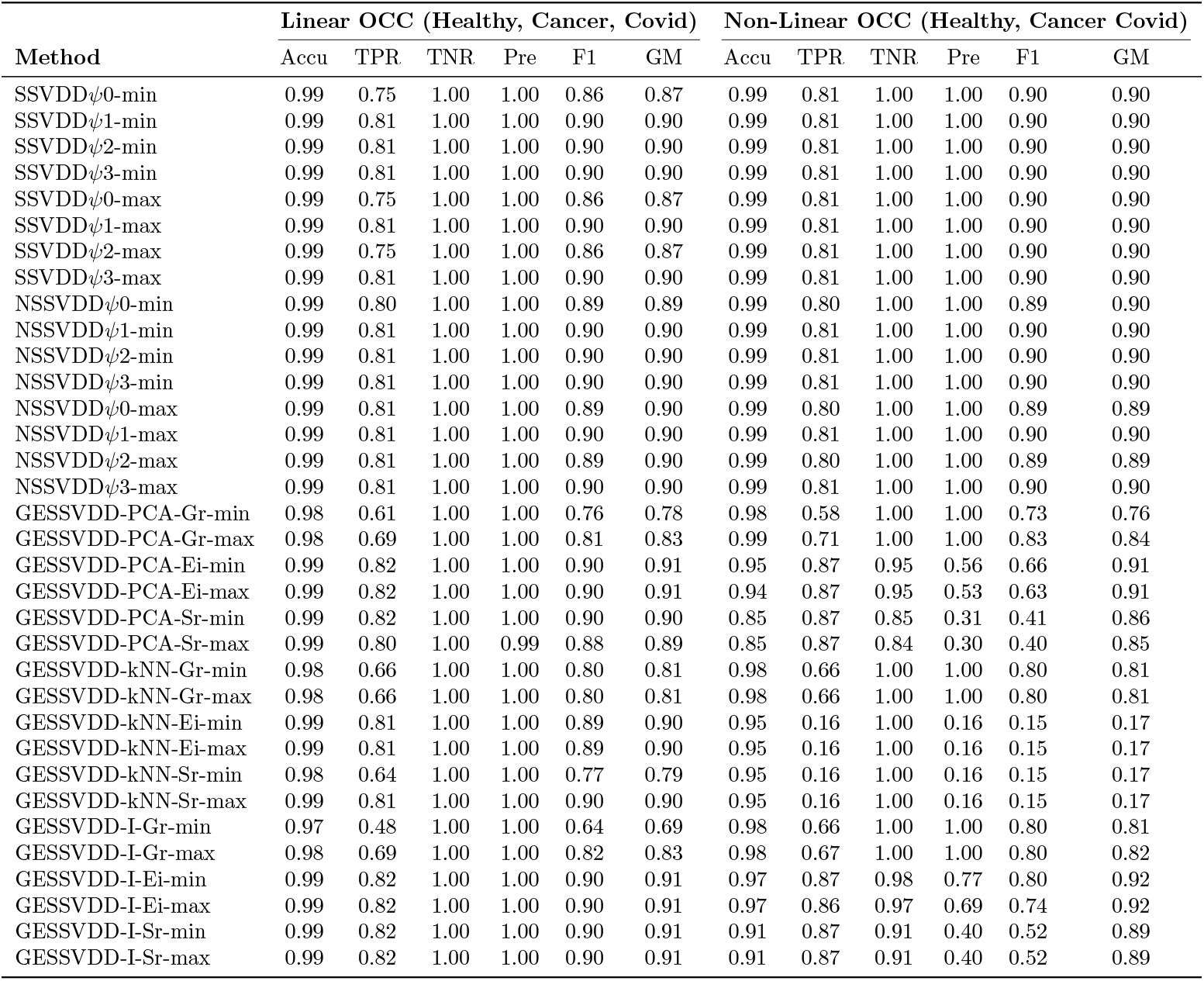
Performance comparison of Linear and Non-Linear OCC methods for (Healthy, Cancer Covid).

## Supplementary 3: Demographic Structure Analysis in the Healthy Cohort

This supplementary presents auxiliary experiments evaluating whether one-class models trained on healthy plasma proteomics profiles encode demographic structure unrelated to disease. Gender-based experiments are reported in Tables 3 and 4, where female and male groups are alternately treated as the target class. Ethnicity-based experiments are reported in Tables 5 and 6, with non-white and white groups treated as the target class, respectively. Across configurations, detection performance remains unstable and near chance level, supporting the conclusion that the learned subspaces do not capture strong demographic stratification.

**Table S3:**
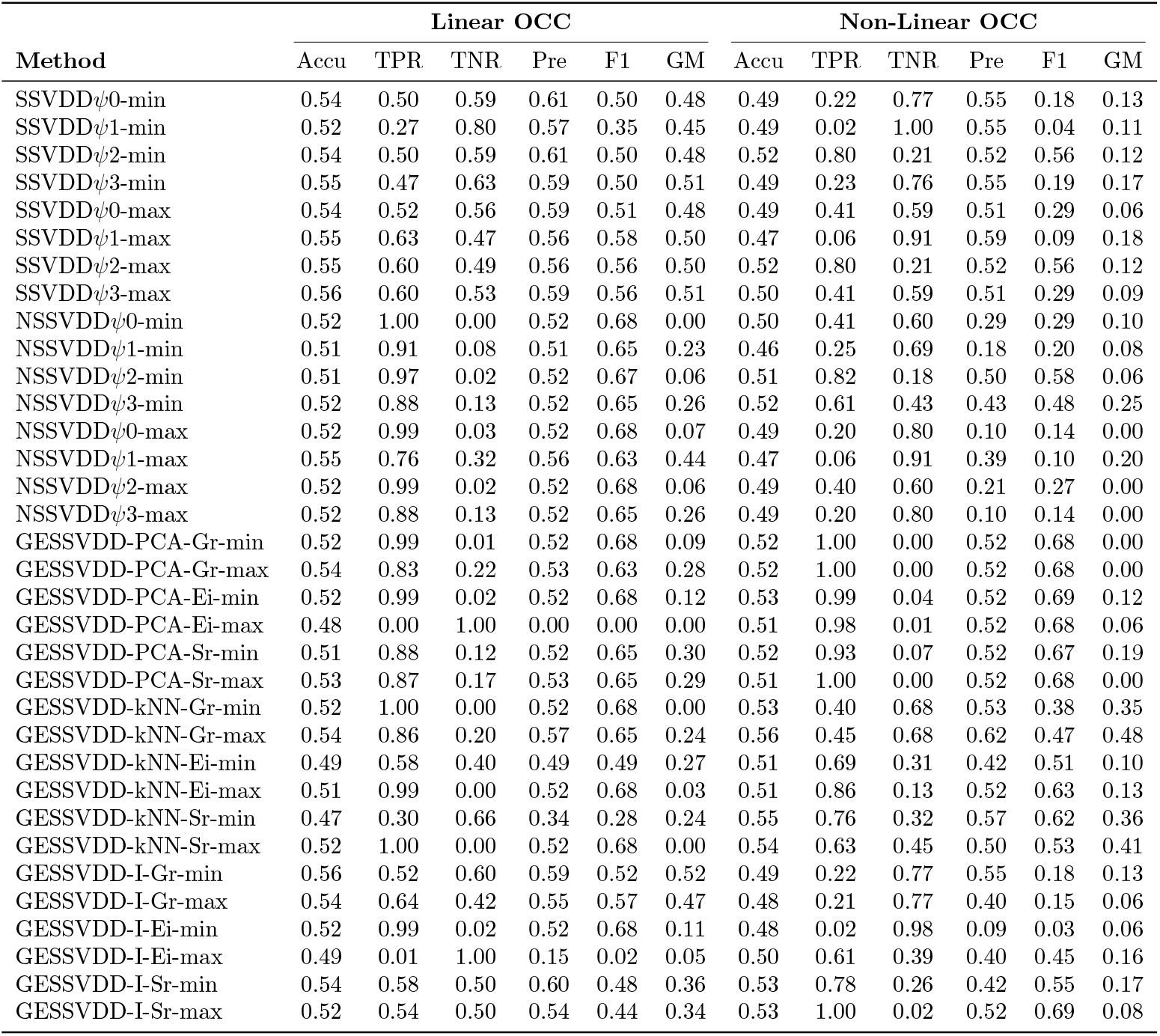
Performance comparison of Linear and Non-Linear OCC methods for Gender Identification in Healthy cohort for target class as FEMALE.

| Method | Linear OCC |  |  |  |  |  | Non-Linear OCC |  |  |  |  |  |
| --- | --- | --- | --- | --- | --- | --- | --- | --- | --- | --- | --- | --- |
|  | Accu | TPR | TNR | Pre | F1 | GM | Accu | TPR | TNR | Pre | F1 | GM |
| SSVDDψ0-min | 0.54 | 0.50 | 0.59 | 0.61 | 0.50 | 0.48 | 0.49 | 0.22 | 0.77 | 0.55 | 0.18 | 0.13 |
| SSVDDψ1-min | 0.52 | 0.27 | 0.80 | 0.57 | 0.35 | 0.45 | 0.49 | 0.02 | 1.00 | 0.55 | 0.04 | 0.11 |
| SSVDDψ2-min | 0.54 | 0.50 | 0.59 | 0.61 | 0.50 | 0.48 | 0.52 | 0.80 | 0.21 | 0.52 | 0.56 | 0.12 |
| SSVDDψ3-min | 0.55 | 0.47 | 0.63 | 0.59 | 0.50 | 0.51 | 0.49 | 0.23 | 0.76 | 0.55 | 0.19 | 0.17 |
| SSVDDψ0-max | 0.54 | 0.52 | 0.56 | 0.59 | 0.51 | 0.48 | 0.49 | 0.41 | 0.59 | 0.51 | 0.29 | 0.06 |
| SSVDDψ1-max | 0.55 | 0.63 | 0.47 | 0.56 | 0.58 | 0.50 | 0.47 | 0.06 | 0.91 | 0.59 | 0.09 | 0.18 |
| SSVDDψ2-max | 0.55 | 0.60 | 0.49 | 0.56 | 0.56 | 0.50 | 0.52 | 0.80 | 0.21 | 0.52 | 0.56 | 0.12 |
| SSVDDψ3-max | 0.56 | 0.60 | 0.53 | 0.59 | 0.56 | 0.51 | 0.50 | 0.41 | 0.59 | 0.51 | 0.29 | 0.09 |
| NSSVDDψ0-min | 0.52 | 1.00 | 0.00 | 0.52 | 0.68 | 0.00 | 0.50 | 0.41 | 0.60 | 0.29 | 0.29 | 0.10 |
| NSSVDDψ1-min | 0.51 | 0.91 | 0.08 | 0.51 | 0.65 | 0.23 | 0.46 | 0.25 | 0.69 | 0.18 | 0.20 | 0.08 |
| NSSVDDψ2-min | 0.51 | 0.97 | 0.02 | 0.52 | 0.67 | 0.06 | 0.51 | 0.82 | 0.18 | 0.50 | 0.58 | 0.06 |
| NSSVDDψ3-min | 0.52 | 0.88 | 0.13 | 0.52 | 0.65 | 0.26 | 0.52 | 0.61 | 0.43 | 0.43 | 0.48 | 0.25 |
| NSSVDDψ0-max | 0.52 | 0.99 | 0.03 | 0.52 | 0.68 | 0.07 | 0.49 | 0.20 | 0.80 | 0.10 | 0.14 | 0.00 |
| NSSVDDψ1-max | 0.55 | 0.76 | 0.32 | 0.56 | 0.63 | 0.44 | 0.47 | 0.06 | 0.91 | 0.39 | 0.10 | 0.20 |
| NSSVDDψ2-max | 0.52 | 0.99 | 0.02 | 0.52 | 0.68 | 0.06 | 0.49 | 0.40 | 0.60 | 0.21 | 0.27 | 0.00 |
| NSSVDDψ3-max | 0.52 | 0.88 | 0.13 | 0.52 | 0.65 | 0.26 | 0.49 | 0.20 | 0.80 | 0.10 | 0.14 | 0.00 |
| GESSVDD-PCA-Gr-min | 0.52 | 0.99 | 0.01 | 0.52 | 0.68 | 0.09 | 0.52 | 1.00 | 0.00 | 0.52 | 0.68 | 0.00 |
| GESSVDD-PCA-Gr-max | 0.54 | 0.83 | 0.22 | 0.53 | 0.63 | 0.28 | 0.52 | 1.00 | 0.00 | 0.52 | 0.68 | 0.00 |
| GESSVDD-PCA-Ei-min | 0.52 | 0.99 | 0.02 | 0.52 | 0.68 | 0.12 | 0.53 | 0.99 | 0.04 | 0.52 | 0.69 | 0.12 |
| GESSVDD-PCA-Ei-max | 0.48 | 0.00 | 1.00 | 0.00 | 0.00 | 0.00 | 0.51 | 0.98 | 0.01 | 0.52 | 0.68 | 0.06 |
| GESSVDD-PCA-Sr-min | 0.51 | 0.88 | 0.12 | 0.52 | 0.65 | 0.30 | 0.52 | 0.93 | 0.07 | 0.52 | 0.67 | 0.19 |
| GESSVDD-PCA-Sr-max | 0.53 | 0.87 | 0.17 | 0.53 | 0.65 | 0.29 | 0.51 | 1.00 | 0.00 | 0.52 | 0.68 | 0.00 |
| GESSVDD-kNN-Gr-min | 0.52 | 1.00 | 0.00 | 0.52 | 0.68 | 0.00 | 0.53 | 0.40 | 0.68 | 0.53 | 0.38 | 0.35 |
| GESSVDD-kNN-Gr-max | 0.54 | 0.86 | 0.20 | 0.57 | 0.65 | 0.24 | 0.56 | 0.45 | 0.68 | 0.62 | 0.47 | 0.48 |
| GESSVDD-kNN-Ei-min | 0.49 | 0.58 | 0.40 | 0.49 | 0.49 | 0.27 | 0.51 | 0.69 | 0.31 | 0.42 | 0.51 | 0.10 |
| GESSVDD-kNN-Ei-max | 0.51 | 0.99 | 0.00 | 0.52 | 0.68 | 0.03 | 0.51 | 0.86 | 0.13 | 0.52 | 0.63 | 0.13 |
| GESSVDD-kNN-Sr-min | 0.47 | 0.30 | 0.66 | 0.34 | 0.28 | 0.24 | 0.55 | 0.76 | 0.32 | 0.57 | 0.62 | 0.36 |
| GESSVDD-kNN-Sr-max | 0.52 | 1.00 | 0.00 | 0.52 | 0.68 | 0.00 | 0.54 | 0.63 | 0.45 | 0.50 | 0.53 | 0.41 |
| GESSVDD-I-Gr-min | 0.56 | 0.52 | 0.60 | 0.59 | 0.52 | 0.52 | 0.49 | 0.22 | 0.77 | 0.55 | 0.18 | 0.13 |
| GESSVDD-I-Gr-max | 0.54 | 0.64 | 0.42 | 0.55 | 0.57 | 0.47 | 0.48 | 0.21 | 0.77 | 0.40 | 0.15 | 0.06 |
| GESSVDD-I-Ei-min | 0.52 | 0.99 | 0.02 | 0.52 | 0.68 | 0.11 | 0.48 | 0.02 | 0.98 | 0.09 | 0.03 | 0.06 |
| GESSVDD-I-Ei-max | 0.49 | 0.01 | 1.00 | 0.15 | 0.02 | 0.05 | 0.50 | 0.61 | 0.39 | 0.40 | 0.45 | 0.16 |
| GESSVDD-I-Sr-min | 0.54 | 0.58 | 0.50 | 0.60 | 0.48 | 0.36 | 0.53 | 0.78 | 0.26 | 0.42 | 0.55 | 0.17 |
| GESSVDD-I-Sr-max | 0.52 | 0.54 | 0.50 | 0.54 | 0.44 | 0.34 | 0.53 | 1.00 | 0.02 | 0.52 | 0.69 | 0.08 |

**Table S4:** Performance comparison of Linear and Non-Linear OCC methods for Gender Identification in Healthy cohort for target class as MALE.

| Method | Linear OCC |  |  |  |  |  | Non-Linear OCC |  |  |  |  |  |
| --- | --- | --- | --- | --- | --- | --- | --- | --- | --- | --- | --- | --- |
|  | Accu | TPR | TNR | Pre | F1 | GM | Accu | TPR | TNR | Pre | F1 | GM |
| SSVDDψ0-min | 0.52 | 0.00 | 1.00 | 0.00 | 0.00 | 0.00 | 0.48 | 1.00 | 0.00 | 0.48 | 0.65 | 0.00 |
| SSVDDψ1-min | 0.52 | 0.00 | 1.00 | 0.20 | 0.01 | 0.03 | 0.48 | 1.00 | 0.00 | 0.48 | 0.65 | 0.00 |
| SSVDDψ2-min | 0.52 | 0.00 | 1.00 | 0.00 | 0.00 | 0.00 | 0.48 | 1.00 | 0.00 | 0.48 | 0.65 | 0.00 |
| SSVDDψ3-min | 0.52 | 0.00 | 1.00 | 0.00 | 0.00 | 0.00 | 0.48 | 1.00 | 0.00 | 0.48 | 0.65 | 0.00 |
| SSVDDψ0-max | 0.51 | 0.00 | 1.00 | 0.00 | 0.00 | 0.00 | 0.48 | 1.00 | 0.00 | 0.48 | 0.65 | 0.00 |
| SSVDDψ1-max | 0.52 | 0.00 | 1.00 | 0.00 | 0.00 | 0.00 | 0.48 | 1.00 | 0.00 | 0.48 | 0.65 | 0.00 |
| SSVDDψ2-max | 0.52 | 0.00 | 1.00 | 0.00 | 0.00 | 0.00 | 0.48 | 1.00 | 0.00 | 0.48 | 0.65 | 0.00 |
| SSVDDψ3-max | 0.51 | 0.00 | 1.00 | 0.00 | 0.00 | 0.00 | 0.48 | 1.00 | 0.00 | 0.48 | 0.65 | 0.00 |
| NSSVDDψ0-min | 0.49 | 0.80 | 0.20 | 0.39 | 0.52 | 0.00 | 0.50 | 0.92 | 0.11 | 0.49 | 0.63 | 0.11 |
| NSSVDDψ1-min | 0.52 | 0.00 | 1.00 | 0.00 | 0.00 | 0.00 | 0.52 | 0.00 | 1.00 | 0.00 | 0.00 | 0.00 |
| NSSVDDψ2-min | 0.46 | 0.64 | 0.28 | 0.45 | 0.53 | 0.42 | 0.48 | 1.00 | 0.00 | 0.48 | 0.65 | 0.00 |
| NSSVDDψ3-min | 0.52 | 0.00 | 1.00 | 0.00 | 0.00 | 0.00 | 0.49 | 0.80 | 0.20 | 0.39 | 0.52 | 0.00 |
| NSSVDDψ0-max | 0.48 | 1.00 | 0.00 | 0.48 | 0.65 | 0.00 | 0.48 | 1.00 | 0.00 | 0.48 | 0.65 | 0.00 |
| NSSVDDψ1-max | 0.52 | 0.00 | 1.00 | 0.10 | 0.01 | 0.03 | 0.52 | 0.00 | 1.00 | 0.20 | 0.01 | 0.03 |
| NSSVDDψ2-max | 0.53 | 0.84 | 0.24 | 0.53 | 0.64 | 0.34 | 0.48 | 1.00 | 0.00 | 0.48 | 0.65 | 0.00 |
| NSSVDDψ3-max | 0.51 | 0.20 | 0.80 | 0.10 | 0.13 | 0.00 | 0.48 | 1.00 | 0.00 | 0.48 | 0.65 | 0.00 |
| GESSVDD-PCA-Gr-min | 0.48 | 1.00 | 0.00 | 0.48 | 0.65 | 0.00 | 0.48 | 1.00 | 0.00 | 0.48 | 0.65 | 0.00 |
| GESSVDD-PCA-Gr-max | 0.53 | 0.48 | 0.58 | 0.57 | 0.45 | 0.45 | 0.48 | 1.00 | 0.00 | 0.48 | 0.65 | 0.00 |
| GESSVDD-PCA-Ei-min | 0.52 | 0.00 | 1.00 | 0.00 | 0.00 | 0.00 | 0.51 | 0.53 | 0.49 | 0.40 | 0.45 | 0.35 |
| GESSVDD-PCA-Ei-max | 0.52 | 0.00 | 1.00 | 0.00 | 0.00 | 0.00 | 0.52 | 0.04 | 0.97 | 0.12 | 0.06 | 0.09 |
| GESSVDD-PCA-Sr-min | 0.52 | 0.00 | 1.00 | 0.00 | 0.00 | 0.00 | 0.53 | 0.30 | 0.73 | 0.37 | 0.31 | 0.35 |
| GESSVDD-PCA-Sr-max | 0.52 | 0.00 | 1.00 | 0.00 | 0.00 | 0.00 | 0.51 | 0.23 | 0.77 | 0.43 | 0.24 | 0.27 |
| GESSVDD-kNN-Gr-min | 0.46 | 0.59 | 0.35 | 0.45 | 0.50 | 0.43 | 0.49 | 0.80 | 0.20 | 0.39 | 0.52 | 0.00 |
| GESSVDD-kNN-Gr-max | 0.53 | 0.31 | 0.74 | 0.50 | 0.36 | 0.43 | 0.48 | 1.00 | 0.00 | 0.48 | 0.65 | 0.00 |
| GESSVDD-kNN-Ei-min | 0.48 | 0.98 | 0.01 | 0.48 | 0.65 | 0.09 | 0.48 | 1.00 | 0.00 | 0.48 | 0.65 | 0.03 |
| GESSVDD-kNN-Ei-max | 0.52 | 0.78 | 0.28 | 0.50 | 0.59 | 0.39 | 0.49 | 0.33 | 0.63 | 0.51 | 0.32 | 0.36 |
| GESSVDD-kNN-Sr-min | 0.48 | 0.96 | 0.04 | 0.48 | 0.64 | 0.13 | 0.47 | 0.91 | 0.06 | 0.47 | 0.62 | 0.11 |
| GESSVDD-kNN-Sr-max | 0.52 | 0.78 | 0.28 | 0.50 | 0.59 | 0.39 | 0.53 | 0.14 | 0.89 | 0.58 | 0.20 | 0.31 |
| GESSVDD-I-Gr-min | 0.52 | 0.00 | 1.00 | 0.00 | 0.00 | 0.00 | 0.48 | 1.00 | 0.00 | 0.48 | 0.65 | 0.00 |
| GESSVDD-I-Gr-max | 0.52 | 0.00 | 1.00 | 0.00 | 0.00 | 0.00 | 0.48 | 1.00 | 0.00 | 0.48 | 0.65 | 0.00 |
| GESSVDD-I-Ei-min | 0.52 | 0.00 | 1.00 | 0.00 | 0.00 | 0.00 | 0.55 | 0.20 | 0.88 | 0.50 | 0.28 | 0.36 |
| GESSVDD-I-Ei-max | 0.52 | 0.00 | 1.00 | 0.00 | 0.00 | 0.00 | 0.51 | 0.02 | 0.97 | 0.08 | 0.03 | 0.06 |
| GESSVDD-I-Sr-min | 0.52 | 0.00 | 1.00 | 0.00 | 0.00 | 0.00 | 0.52 | 0.01 | 1.00 | 0.30 | 0.02 | 0.06 |
| GESSVDD-I-Sr-max | 0.52 | 0.00 | 1.00 | 0.00 | 0.00 | 0.00 | 0.53 | 0.29 | 0.76 | 0.34 | 0.26 | 0.24 |

**Table S5:**
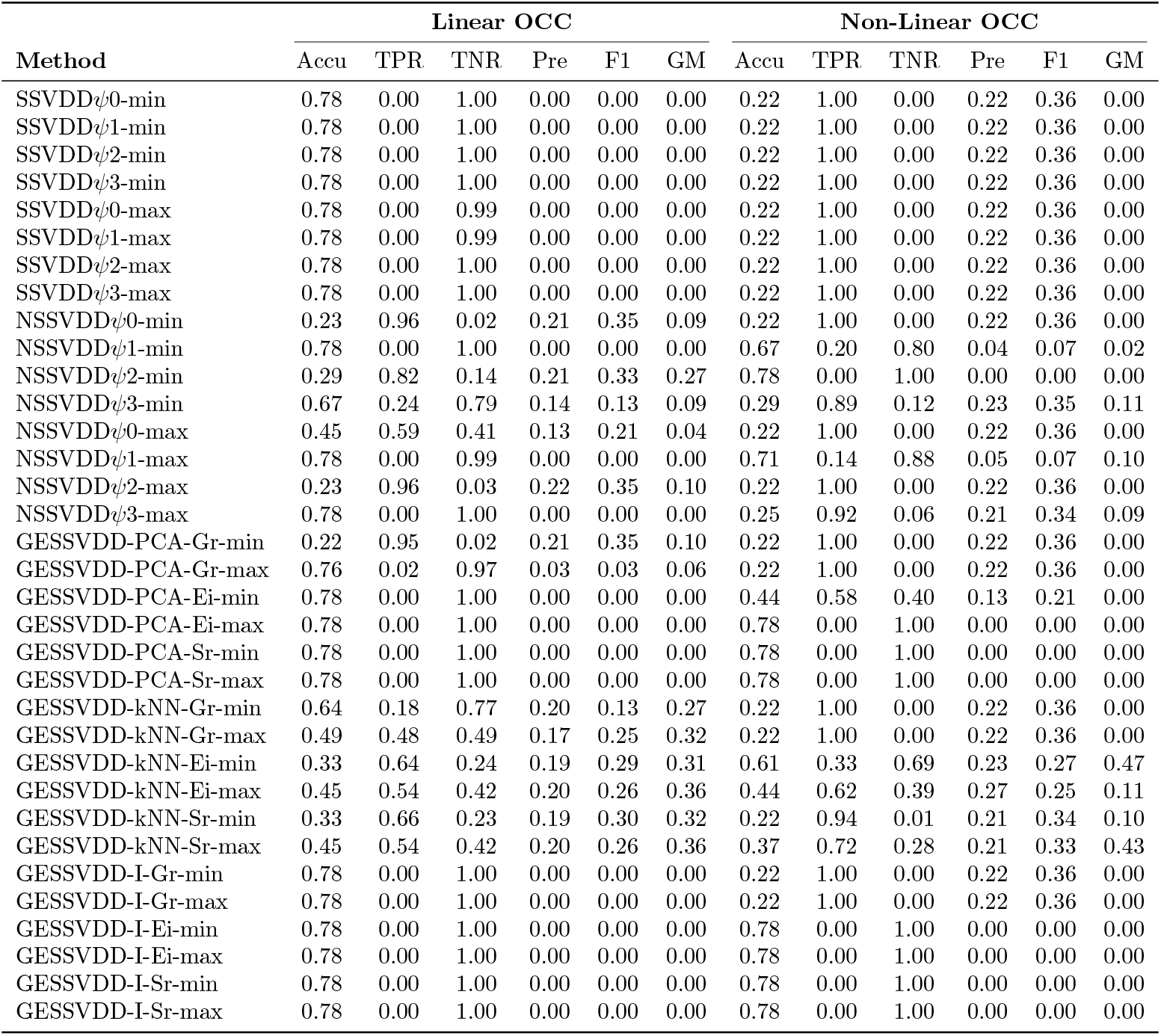
Performance comparison of Linear and Non-Linear OCC methods for Ethnicity target class as NON WHITE.

**Table S6:**
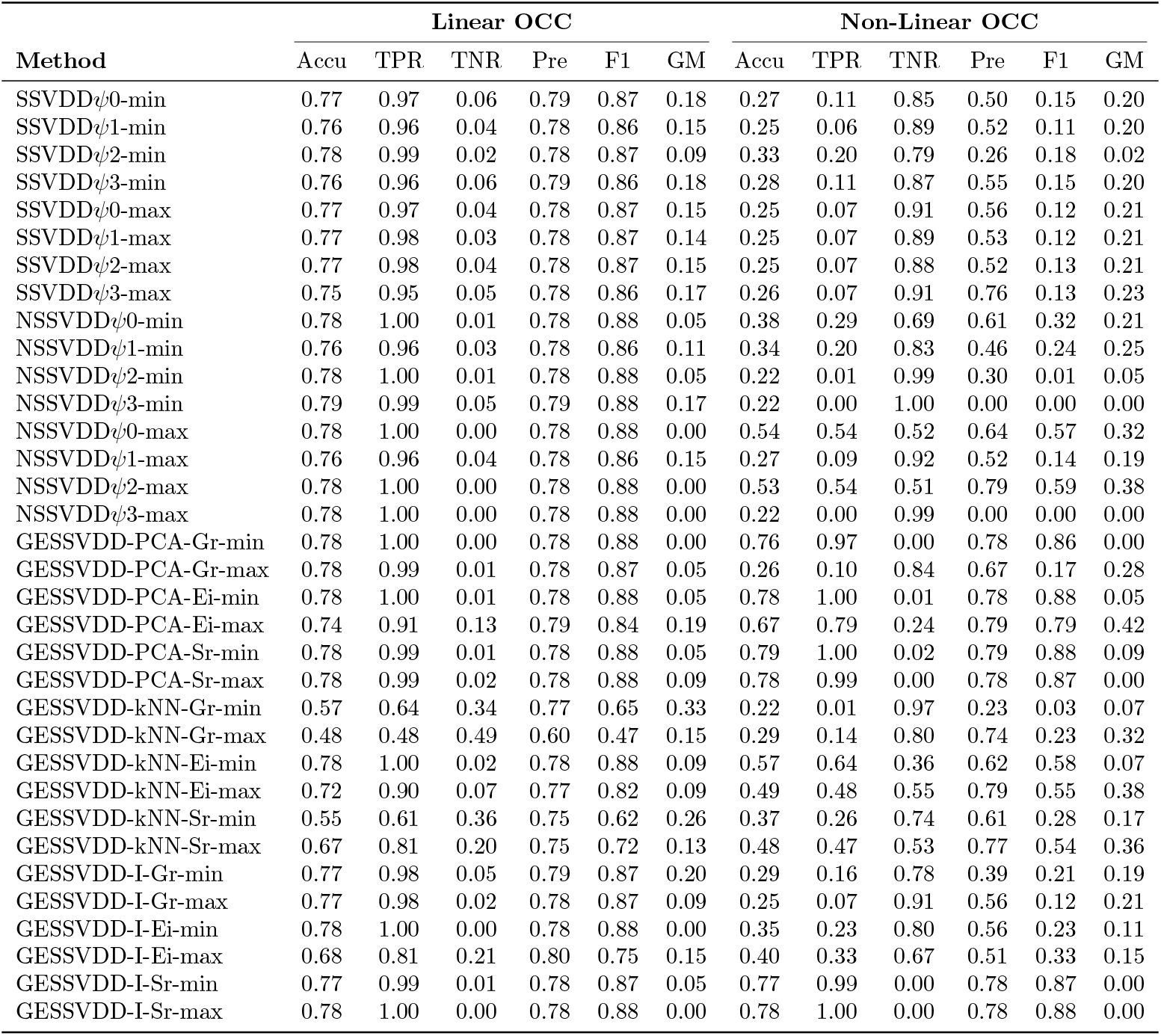
Performance comparison of Linear and Non-Linear OCC methods for Ethnicity target class as WHITE.

## Supplementary 4: Dataset Composition and PCA Visualization

**Fig. S1:**
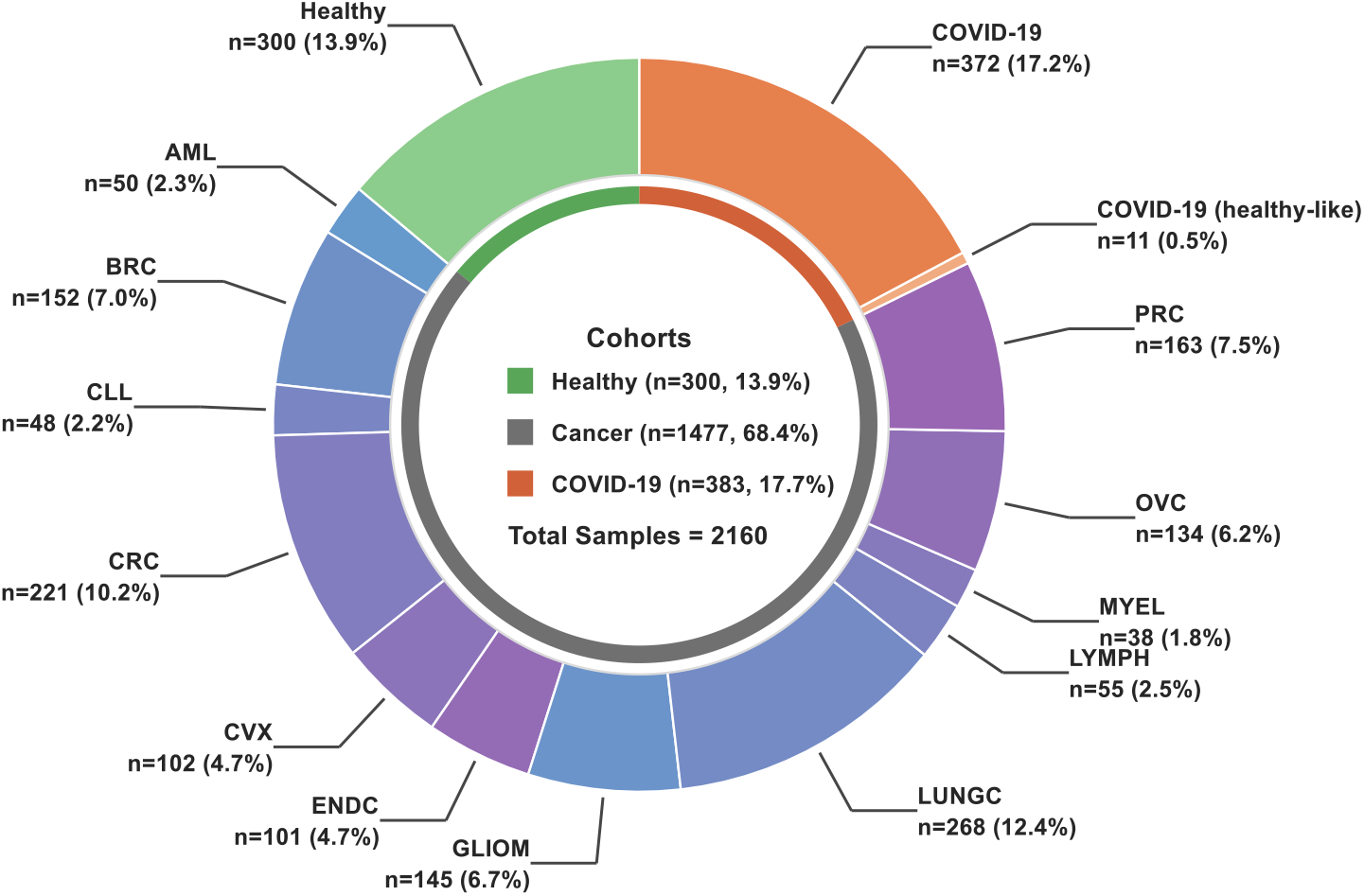
Composition of the plasma proteomics dataset across cohorts. Pie chart showing the number and proportion of samples from the healthy reference cohort, multiple cancer types, and an independent COVID-19 cohort. Individual cancer types are shown separately, while COVID-19 samples include both COVID-positive and healthy-like individuals within the COVID-19 cohort. Percentages indicate the fraction of the total dataset represented by each subgroup.

**Fig. S2:**
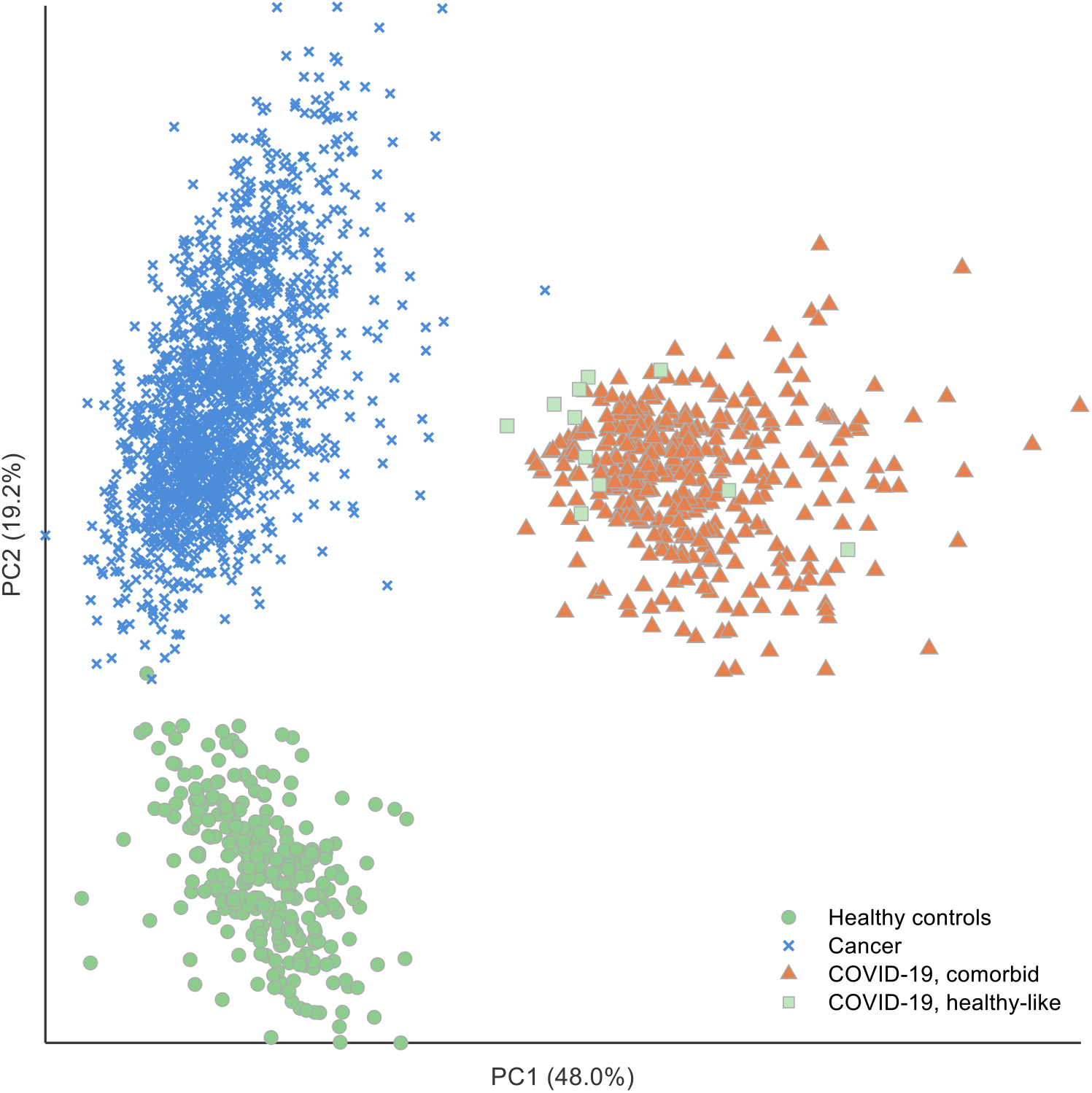
Principal component analysis of plasma proteomic profiles across cohorts. Two-dimensional projection of plasma proteomic samples onto the first two principal components (PC1 and PC2), computed from a curated set of 62 shared protein assays measured using Olink Proximity Extension Assay. The analysis includes samples from a healthy control cohort, multiple cancer cohorts, and an independent COVID-19 cohort. Prior to PCA, protein expression values were globally z-score normalized across all samples. Each point represents an individual sample, with colors and markers indicating cohort membership: healthy controls, cancer patients, COVID-19 healthy-like individuals, and COVID-19 individuals with comorbidities. The plot provides an exploratory visualization of the global variance structure of plasma proteomic profiles across health and disease.

## Footnotes

1 codes will be made public upon final acceptance of paper.

